# Predictive acoustical processing in human cortical layers

**DOI:** 10.1101/2025.01.09.632099

**Authors:** Lonike K. Faes, Isma Zulfiqar, Luca Vizioli, Zidan Yu, Yuan-Hao Wu, Jiyun Shin, Martijn A. Cloos, Ryszard Auksztulewicz, Lucia Melloni, Kamil Uludag, Essa Yacoub, Federico De Martino

## Abstract

In our dynamic environments, predictive processing is vital for auditory perception and its associated behaviors. Predictive coding formalizes inferential processes by implementing them as information exchange across cortical layers and areas. With laminar-specific blood oxygenation level dependent we measured responses to a cascading oddball paradigm, to ground predictive auditory processes on the mesoscopic human cortical architecture. We show that the violation of predictions are potentially hierarchically organized and associated with responses in superficial layers of the planum polare and middle layers of the lateral temporal cortex. Moreover, we relate the updating of the brain’s internal model to changes in deep layers. Using a modeling approach, we derive putative changes in neural dynamics while accounting for draining effects. Our results support the role of temporal cortical architecture in the implementation of predictive coding and highlight the ability of laminar fMRI to investigate mesoscopic processes in a large extent of temporal areas.

## Introduction

The ability to form predictions about incoming sensory stimuli is crucial for everyday behavior. Sounds that reach our ears are often noisy and incomplete and by anticipating what we will hear next, the brain can efficiently filter and prioritize behaviorally relevant sounds, facilitating for example auditory stream segregation and understanding speech in noise [1–3]. Aberrant predictive processing is even hypothesized to underlie phantom perception such as tinnitus [4] or hallucinations [5–7]. A widely referenced theory of perceptual processing is predictive coding [8,PC - 9], suggesting that this process of inference occurs through hierarchical exchange of information across brain areas, attributing specialized roles to feedforward and feedback streams by capitalizing on the known anatomical separation across cortical layers [10,but see 11]. In PC, descending feedback is hypothesized to convey predictions of lower-level representations that are compared to sensory input at lower stages of the hierarchy. Conversely, instead of forwarding the sensory input, the discrepancy between the prediction and this sensory input (i.e. the prediction-error) is fed forward to higher-order brain areas [12–14].

Evidence in favor of distinct computational roles of cortical layers for PC mainly comes from invasive animal studies [15–20]. Nevertheless, it remains an open question whether this version of PC is translatable to humans [21]. Invasive electrophysiological measures in patients offer a potential means of bridging the gap between animal and human literature. Although direct recordings using laminar electrodes in humans are feasible, to date, these electrodes still offer recordings from a limited number of areas across the cortex and have not yet been used to test test hypotheses related to PC [22–24]. Ultra-high field functional magnetic resonance imaging (UHF fMRI) offers an alternative to non-invasively investigate the laminar flow of information in the human brain. The use of UHF fMRI has yielded insights into the specific contributions of cortical layers in predictive processing. A wealth of studies has centered on the visual [25–28] and the somatosensory cortex [29] and have broadly confirmed the hypotheses regarding the mesoscopic implementation of PC. Predictive coding has been proposed as a general framework for understanding how perceptual processing occurs in the brain. As such, it is commonly assumed that mechanistic implementations generalize across senses; yet its implementation in the human auditory cortex remains untested. Moreover, even though the existence of hierarchy is an important assumption of PC [for a review see e.g., 13] and has been explored in both rodents [17,30], non-human primates [15,20,31,32] and humans [33], the laminar profile of hierarchical information exchange also remains largely unexplored in humans [but see 26].

In this work, we present the first human laminar fMRI results investigating how the violation of expectation modulates responses in multiple regions of the auditory cortex probed by a cascading oddball paradigm. To this end, we collected high spatial resolution (0.8 mm isotropic) fMRI data using gradient-echo blood oxygenation level-dependent (GE-BOLD). The data was analyzed using a previously proposed laminar BOLD model [34], to disentangle underlying neuronal activity from (unrelated) vascular changes within the dynamic causal modeling (DCM) framework [35]. We show distinct roles of cortical laminae in processing unpredictable and mispredicted tone sequences compared to predictable sequences. More specifically, for prediction violations, we find error-related signals in superficial layers in a lower-level auditory region, whereas we find middle layer modulation in a higher-order temporal area. Furthermore, a widespread deep layer effect is present across the temporal cortex, possibly indicating global updating of the internal model of the brain both in mispredicted and unpredictable tone sequences.

## Results

### Task and fMRI

Ten healthy volunteers passively listened to a sequence of four tones. Their local and global predictability was manipulated while the BOLD response was measured across multiple hierarchical auditory areas (Figure 2A and 2C) at submillimeter resolution (0.8 mm isotropic) with gradient-echo echo-planar-imaging (GE-EPI). The first three ‘contextual’ frequency tones (493.9, 659.3, and 987.8 Hz) were common to all sequences, and could either be ascending (3 sequences) or descending (3 sequences). We expected that the three successive ascending or descending tones would create a strong expectation, while minimizing local adaptation driven by e.g. the repetition of the same tone as in classical oddball paradigms [13]. The fourth tone in the sequences could either fulfill (*predictable* condition) or violate (*mispredicted* condition) these expectations (see Figure 1A). Finally, in a third condition (*unpredictable*), the sequence of the first three tones was rearranged (see Figure 1A) to reduce the contextual expectation of the last tone, thereby reducing local predictability of the fourth tone and designed to result in a reduced magnitude of local error signals.

**Figure 1.**
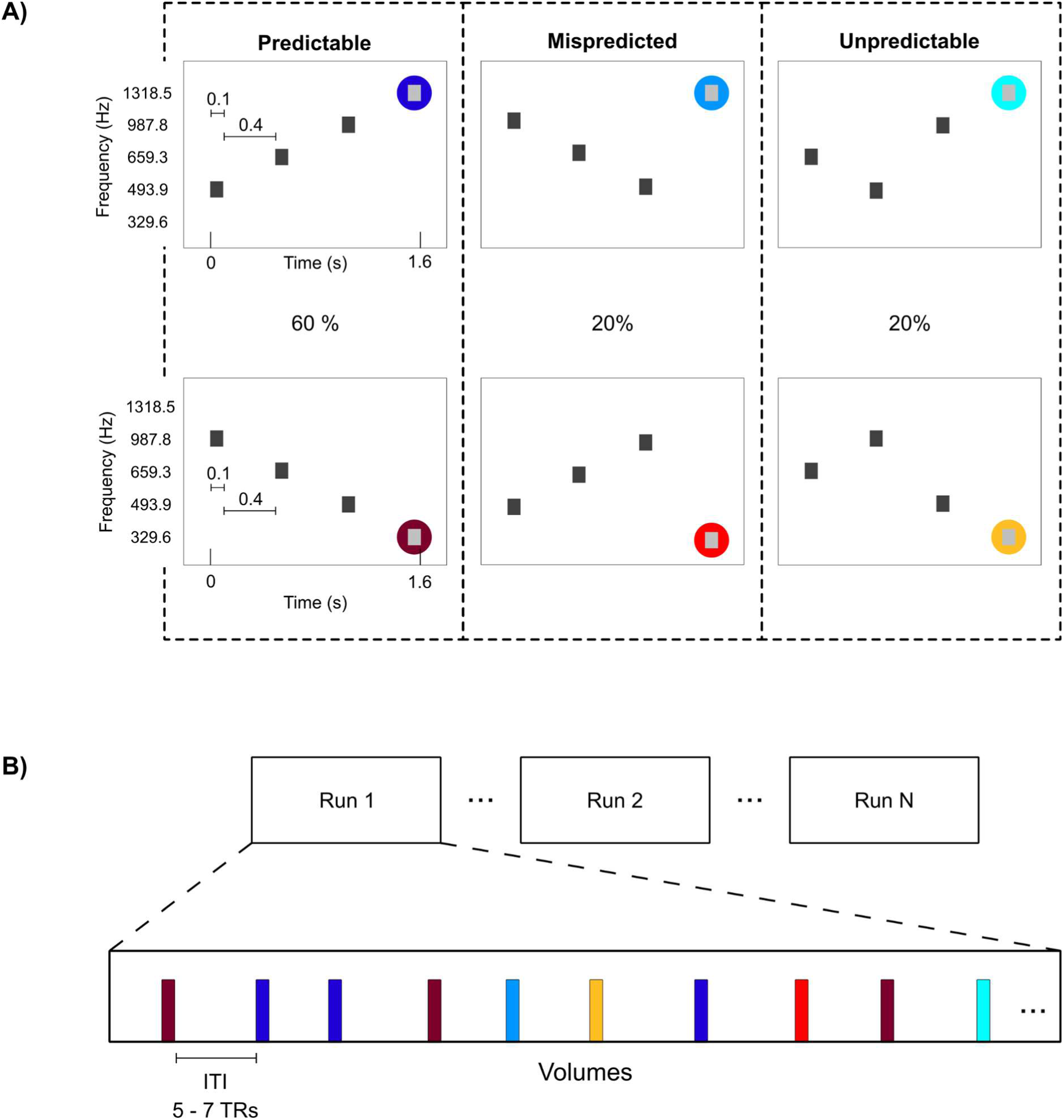
Timing and conditions of the experimental paradigm. A) Three types of conditions were presented. The first three tones either followed a sequence (ascending/descending) that elicited a strong prediction (predictable and mispredicted) or the order of the sequence was scrambled, decreasing the predictability of the last tone (unpredictable). The fourth tone could either fulfill the prediction (predictable) or violate it (mispredicted), creating a local expectation. Ascending or descending sequences could end in a low or high frequency tone. Across the experiment, predictable conditions were presented more often than the mispredicted and unpredictable conditions (60-20-20) creating a global expectation that the Predictable condition was more likely. B) Each session consisted of 6 to 8 runs. In each run, 36 trials were presented in a slow event-related design with an inter-trial interval of 5 to 7 TRs. The order of conditions was randomized.

**Figure 2.**
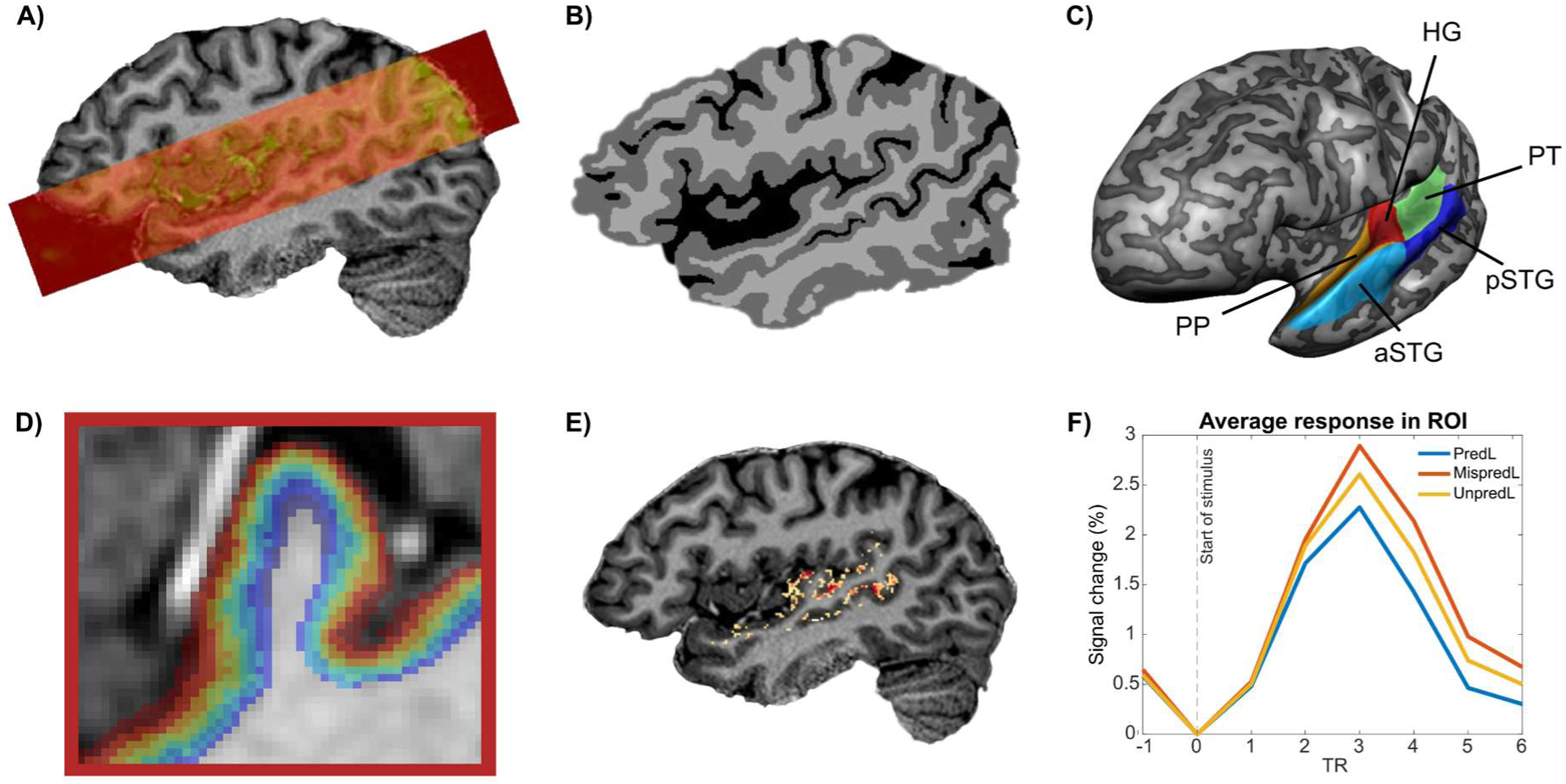
Preprocessing before modeling. A) Coverage of the functional slab overlayed on the high-resolution anatomy. B) Manually edited segmentation of white matter, gray matter and cerebral spinal fluid. C) Example of the regions of interest drawn on a slightly inflated mid gray matter surface. HG = Heschl’s gyrus, PP = planum polare, PT = planum temporale, aSTG = anterior superior temporal gyrus, pSTG = posterior superior temporal gyrus. D) The gray matter curvature was divided into equivolume layers. Illustration of the depth estimates of Heschl’s Gyrus in one volunteer. E) We then selected voxels that responded to auditory stimulation at p<0.05 uncorrected. F) Per ROI, we average the response across trials of each condition. Here, we average all trials of the conditions ending in a low frequency (PredL, MispredL and UnpredL).

Moreover, local predictability (within a sequence) was manipulated alongside global predictability (within the experiment). To induce the latter, the predictable condition was presented more frequently (60%, 20%, and 20% respectively for the three condition-types, see Figure 1A). As a result, we expected global error signals to be present in both mispredicted and unpredictable responses while local errors would be reduced for unpredictable conditions. Each session consisted of 6 to 8 runs. Within each run, the conditions were randomized and presented using a slow event-related design with an inter-trial interval ranging between 8 to 11.2 seconds (5 to 7 TRs - Figure 1B).

### Submillimeter cortical auditory responses

Our functional data slab encompassed the bilateral auditory cortex (Figure 2A). The anatomical data (0.75 mm isotropic) were first upsampled (0.4 mm isotropic) and then automatically segmented into white matter (WM), gray matter (GM) and cerebral spinal fluid (CSF) and manually corrected to accurately delineate the gray matter ribbon in the bilateral temporal cortices (Figure 2B). Using cortical depth estimation, a mid-gray matter surface was created from this high-resolution segmentation, on which we defined five regions of interest (ROIs) based on macro-anatomical landmarks [36 - Figure 2C]. Per ROI, the cortical depth of every voxel together with the local curvature information led to the assignment of each voxel to different cortical depths using the equivolume approach (Figure 2D). First, we localized the voxels responding to sounds (all conditions versus baseline, t >= 2.0, p<0.05 uncorrected; Figure 2E). Subsequently, the analysis proceeded with generating (in each individual subject data and ROI layer) event-related averages (ERA) per condition. For illustration purposes, figure 2F shows the ERA of one of the ROIs in a single volunteer, averaged across all layers. See Figure S1 for activation maps, parcellation of ROIs and an illustration of the draining effect in individual subjects.

GE-BOLD acquisitions are commonly used to survey the cortex at laminar resolution due to its high functional contrast to noise ratio (fCNR) and large coverage [37,38]. This large coverage is especially beneficial for investigating the auditory cortex, since the temporal lobe can be imaged bilaterally [39]. Yet, a downside of GE-BOLD data is its higher sensitivity to (unwanted) macrovasculature effects [e.g. draining veins - 40]. To mitigate this effect, we have applied a validated [41] generative laminar BOLD model [34], incorporated within the DCM framework allowing the estimation of putative underlying laminar neuronal dynamics [see e.g., 35 and Figure 3A]. Using this generative model, we compared the response of mispredicted and unpredictable conditions to the response of predictable ones (in every participant and ROI). In particular, the driving input (i.e. a sequence of four tones) present in all conditions was accompanied by an additional modulatory input (see Figure 3B). In the mispredicted condition, this entailed that modulatory input occurred at the time point at which the prediction was violated (i.e. when the fourth tone was presented). In the unpredictable condition, the modulatory input was modeled at the third and fourth tones to capture the modulatory effect both at the moment at which the context was disrupted (i.e. the first two tones follow a different pattern than the last two tones) and at the end of the sequence to capture the effects of contextual disruption on the processing of the fourth tone.

**Figure 3.**
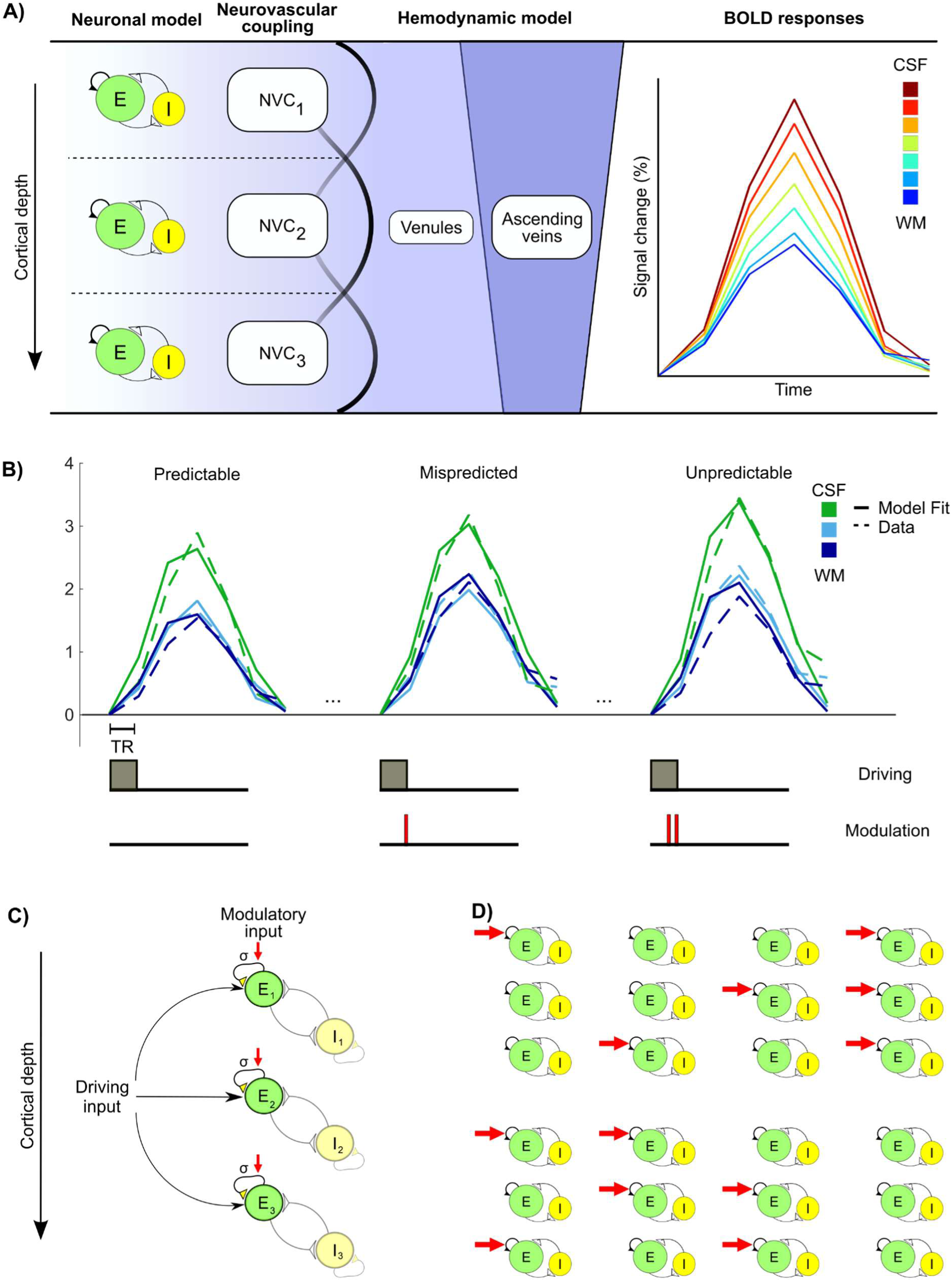
Modeling. A) Schematic of the biophysical model [adapted from 35]. We fixed the number of neuronal depths to 3 and modeled the data at several vascular depths. NVC = neurovascular coupling. B) Example event-related averages of both the data and model fit for three conditions (top panel) visualized for 3 depths; and the timing at which the driving input and the modulatory input come in (bottom panel). C) Schematic of the neuronal model [adapted from 35] to illustrate the target of the driving and modulatory input. D) A total of 8 models were generated, illustrating the distinct layers that are targeted by modulatory input.

The target of this modulatory input was the self-excitation connection of the excitatory neuronal population as this has been previously shown to modulate changes in the overall amplitude of the BOLD response [Figure 3C - and see 35]. Moreover, in electrocorticography (ECoG) data the self-excitatory parameter has been linked to gain modulation in auditory areas when content predictability was manipulated [42]. Note that due to the relatively short inter-stimulus interval, we only modeled the effect of the modulation of self-excitation and refrained from modeling the BOLD undershoot as we did not capture a sufficient number of time points to model it. Within the DCM framework, we compared [using Free Energy - 43] eight models, each hypothesizing distinct neuronal layer(s) targeted by the modulatory input (Figure 3D). Apart from the winning model (see Figure S2), this approach allowed us to estimate (in each participant and ROI) the strength of the modulatory input across the three neuronal depths induced by mispredicted and unpredictable conditions (see Figure S3 for data without modeling).

### Error signaling and model updating for mispredicted stimuli

We first investigated how mispredicted stimuli are represented in the mesoscopic cortical architecture compared to predictable ones. We hypothesized this comparison to highlight the strongest violation effects as well as global update of the internal model (in response to the representation of a rare event). To this end, we averaged across sequence-type (i.e. ascending or descending sequences) and hemispheres. Figure 4 illustrates (across all subjects and ROIs) the layer-dependent modulation in self-recurrent excitation that best explains the difference between mispredicted and predictable conditions. Our results demonstrate a distinct role of cortical depths in processing the violation induced by mispredicted sequences. In particular, we observed a significant effect in deep layers in all five ROIs (HG: t(9) = 4.88, p_FDR-corr_ = 0.008; PP: t(9) = 6.53, p_FDR-corr_ = 0.005; PT: t(9) = 4.37, p_FDR-corr_ = 0.005; aSTG: t(9) = 6.90, p_FDR-corr_ = 0.005; pSTG: t(9) = 5.88, p_FDR-corr_ = 0.005). Interestingly, we only observed a selective effect in the superficial layers in planum polare (t(9) = 3.75, p_FDR-corr_ = 0.005) which we did not observe in any other ROI. Furthermore, in middle layers a significant effect was present solely in the posterior superior temporal gyrus (t(9) = 4.69, p_FDR-corr_ = 0.005), and was not significant in the other ROIs. See table S1 for an overview of the t- and p-values of all ROIs and layers, including the ones that did not reach significance. Statistical significance was assessed by a non-parametric t-test (i.e. the null distribution was estimated using permutations obtained by sign flipping the modulation values) and was corrected for multiple comparisons using false discovery rate (FDR; across regions and layers). For the results prior to modeling, see Figure S3B.

**Figure 4.**
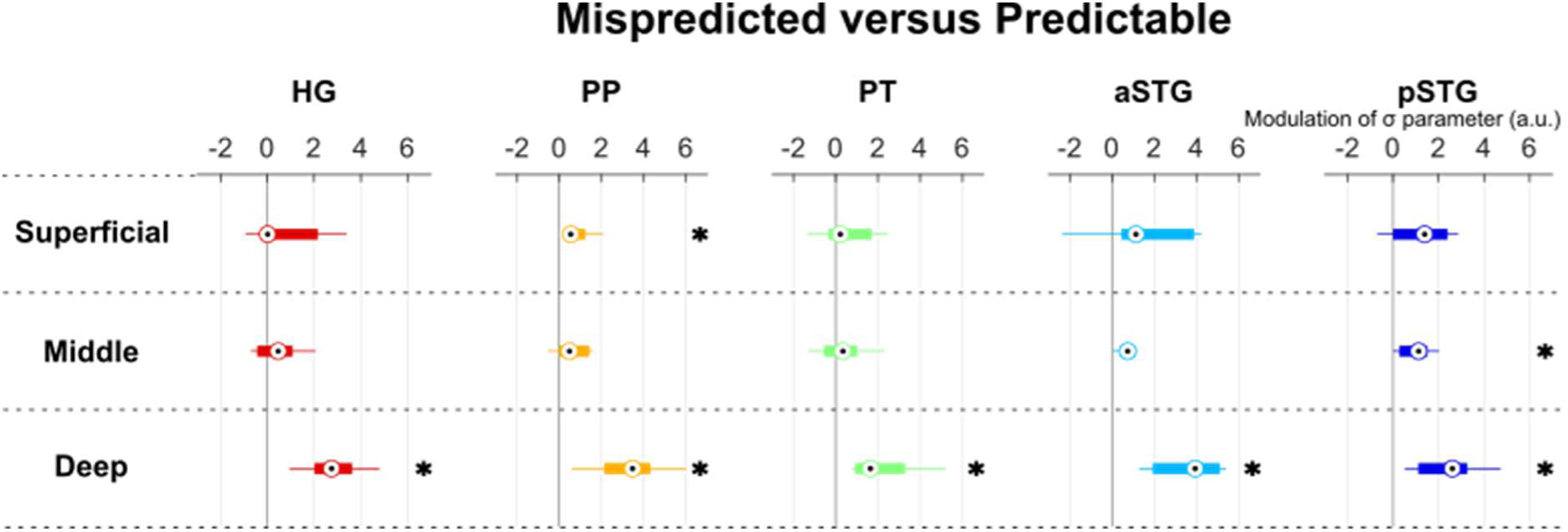
Neuronal modulation of mispredicted stimuli compared to predictable stimuli. For each ROI and cortical layers, the modulation values of the σ parameter (a.u.) that best explains the difference between mispredicted and predictable sequences is plotted across participants. Dots in circles indicate the median value and bars illustrate the interquartiles range (IQR) across participants, with whiskers extending for one time the IQR beyond the limits of the IQR (roughly representing 83% of the data). All ROIs exhibit significant modulation in the deep layers in response to mispredicted stimuli; while a superficial layer effect was present only in planum polare and a middle layer effect was visible only in posterior superior temporal gyrus. HG = Heschl’s gyrus, PP = planum polare, PT = planum temporale, aSTG = anterior superior temporal gyrus, pSTG = posterior superior temporal gyrus. * indicates *p*_FDR-corr_ < 0.05.

### Processing of unpredictable sequences

We then assessed the modulation induced by unpredictable sequences (compared to predictable ones). We hypothesized that the unpredictable sequences would have reduced expectations for the last tone because of the scrambled order of the contextual ones and thus reduced (local) errors, while preserving the need for a global update of the internal model because of the rarity of the events. Using a similar procedure as for the analysis of modulation induced by mispredicted stimuli (averaging across hemisphere and sequence-type, significance testing using permutations for significance testing and FDR corrections for multiple comparisons), we observed significant effects in deep layers in HG (t(9) = 3.41, p_FDR-corr_ = 0.049), PP (t(9) = 3.57, p_FDR-corr_ = 0.049) and aSTG (t(9) = 4.70, p_FDR-corr_ = 0.049), but not in PT and pSTG. In the middle and superficial layers, we found no evidence for a condition-specific modulation (Figure 5). See table S2 for an overview of the t- and p-values of all ROIs and layers, including the ones that did not reach significance. For the results prior to modeling, see Figure S3B.

**Figure 5.**
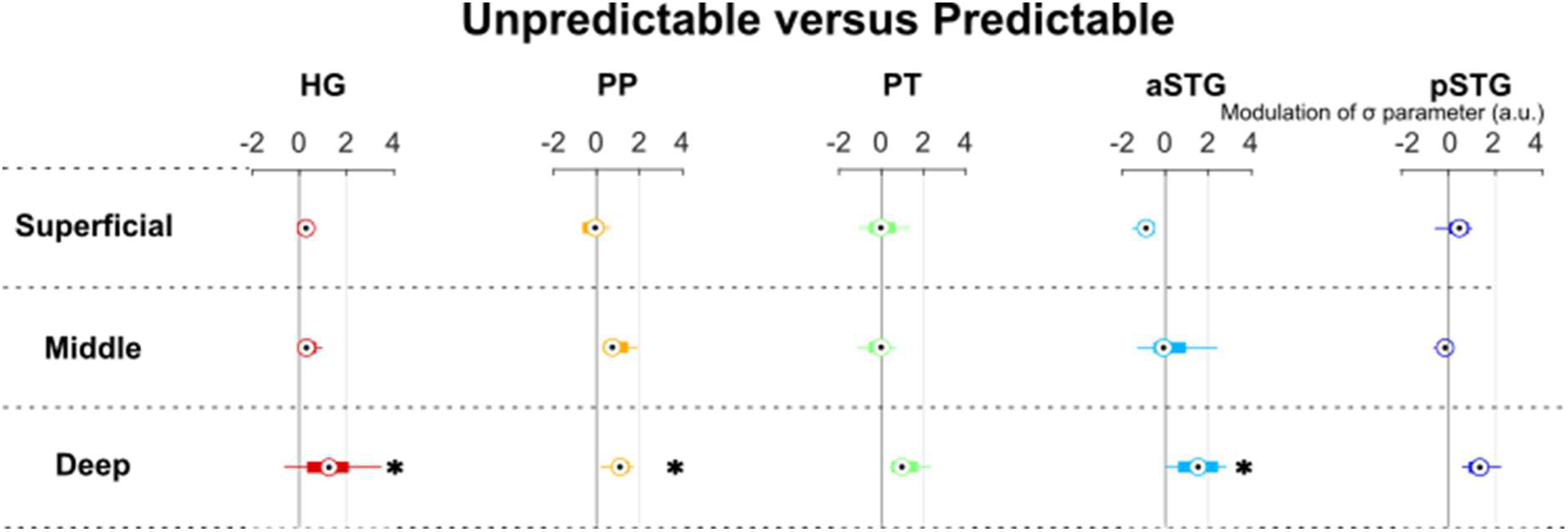
Neuronal modulation of unpredictable stimuli compared to predictable stimuli. A condition-specific modulation of the σ parameter (a.u.) is present in deep layers in Heschl’s gyrus, planum polare and anterior superior temporal gyrus. This figure illustrates the modulation values of sigma that best explains the difference between unpredictable and predictable sequences across participants. Dots in circles indicate the median value and bars illustrate the interquartiles range (IQR) across participants, with whiskers extending for one time the IQR beyond the limits of the IQR (roughly representing 83% of the data). There is no evidence of modulation in the middle and superficial layers. HG = Heschl’s gyrus, PP = planum polare, PT = planum temporale, aSTG = anterior superior temporal gyrus, pSTG = posterior superior temporal gyrus. * indicates p < 0.05.

## Discussion

To map the mesoscopic circuitry relevant for processing of the violation of predictions in the auditory cortex, we measured cortical depth dependent responses to stimuli that adhered to, reduced or violated expectations. Departing from previous research, to maximize neuronal specificity, we employed a modeling approach to account for venous drainage in the most commonly used laminar fMRI acquisition (i.e. GE-EPI). The use of GE-EPI allowed us to explore the hierarchical organization of layer-specific signals across a large extent of the temporal lobe. Our findings revealed modulation in superficial layers in planum polare and in the middle layers of the posterior STG in response to the violation of the internal model. Moreover, we found consistent modulation in deep layers in all our ROIs when contrasting predictable to mispredicted stimuli. In contrast, the unpredictable stimuli modulated the response in deep layers in some of the temporal regions, but did not elicit a notable modulation in the superficial or middle layers.

We demonstrated that mispredicted sounds, which violated both the local predictive context and global expectations, elicit a significant modulation in the superficial layers of PP. The involvement of superficial layers to the processing of errors has been hypothesized by the proposed microcircuit for predictive coding [12]. Evidence in favor of this hypothesis so-far has been gathered by invasive research in animals, that has identified superficial layer activity associated with prediction errors in the visual cortex of mice (Gallimore et al., 2023; Hamm et al., 2021; Jordan & Keller, 2020; G. B. Keller et al., 2012). In the auditory modality, invasive recordings in non-human primates have also identified supragranular layers of the primary auditory cortex to be responsive to the violation of predictions induced by the repetition of the same tone and disentangled from stimulus specific adaptation [20]. In humans, the relevance of the mesoscopic architecture to the processing of errors is so-far supported by a study conducted in the visual domain demonstrating that deviant stimuli are only decodable in superficial layers of V1 [28]. Our results extend this research and provide, for the first time in the human auditory cortex, evidence of the involvement of superficial cortical layers in processing prediction errors. In our data errors are signaled first by superficial layers of PP, indicating that the more complex acoustic rule (compared to repeating the same tone) of our cascade stimuli (minimizing stimulus specific adaptation) may move the relevant signaling at a higher hierarchical level compared to previous research in non-human primates [20].

Interestingly, the larger coverage afforded by UHF laminar fMRI also highlighted a modulation of the middle layers of pSTG in response to prediction violations. Within the predictive coding (PC) framework, middle cortical layers at higher hierarchical levels receive errors fed forward by preceding processing stages, suggesting routing of information from PP to pSTG in response to the violation of predictions. The hierarchical nature of predictions and prediction errors has been previously reported in rodents [30], non-human primates [20,31,32] and humans [33], while the relevance of superficial and deep layers for errors and predictions respectively has been evidenced in non-human primates [15]. A previous study using somatosensory stimulation in humans has identified that middle layer activity is driven by predictable sensory stimuli in the primary somatosensory cortex [see e.g., 29]. Here we highlight the distinct relevance of superficial and middle layers in the hierarchical organization of cortical error processing. This hypothesized difference in hierarchical level of PP and pSTG is supported considering the alignment of the architectonic model developed in non-human primates to the human temporal cortex [see e.g., 44]. PP is considered to be the human homologue of the belt middle medial (MM) and rostro medial (RM) areas [45–47] while the pSTG is thought to be the homologue to the caudal parabelt (CP). In non-human primates, these areas have been shown to be connected, suggesting a similar connectivity in humans [48]. It is thus conceivable that error signals could travel from superficial layers of PP to middle layers of pSTG (see Figure 6A and 6B).

**Figure 6.**
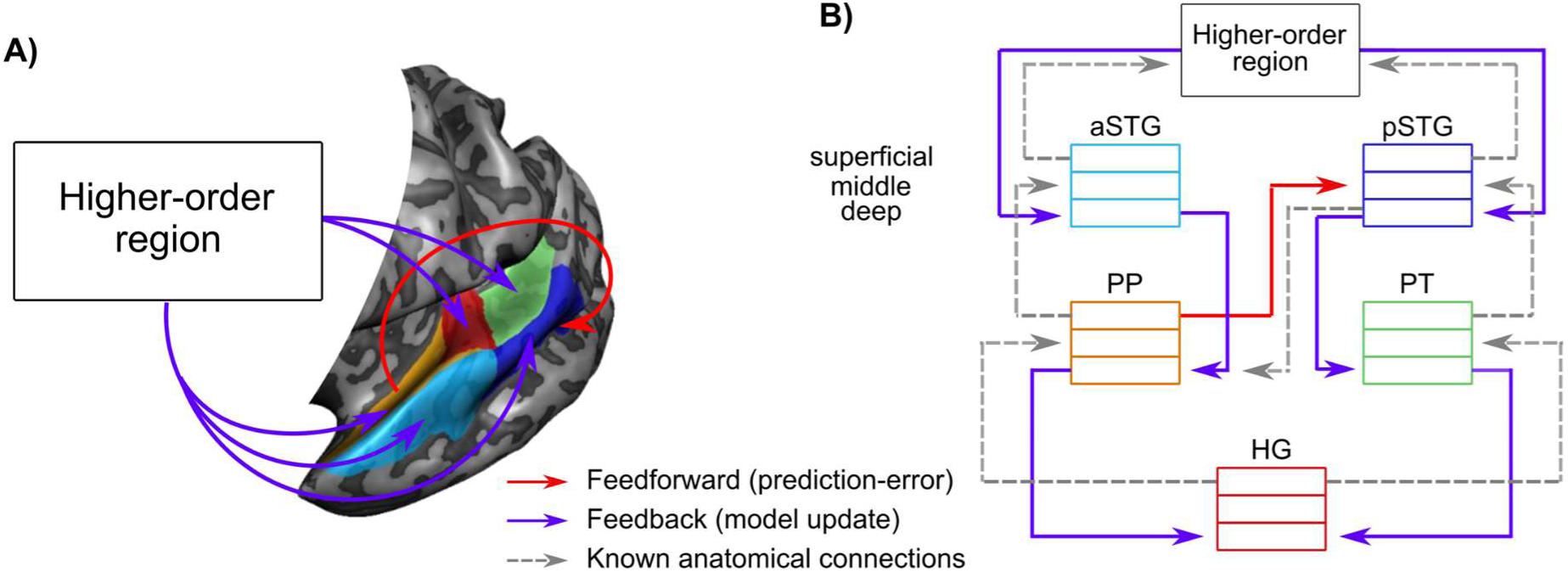
Overview of hypothesized information exchange across the auditory cortical hierarchy. A) Schematic overview of hypothesized feedforward and feedback streams across different cortical areas. Feedforward streams are thought to convey a prediction-error, whereas the feedback information represents a global model update signal. B) Schematic overview of the hypothesized laminar information exchange. Feedforward prediction-error is conveyed from superficial layers in planum polare to the middle layer of posterior superior temporal gyrus. The global model update signals are conveyed to deep layers of all areas. Gray arrows indicated the other anatomical connections described in previous research [48,53]. HG = Heschl’s gyrus, PP = planum polare, PT = planum temporale, aSTG = anterior superior temporal gyrus, pSTG = posterior superior temporal gyrus.

We also found a robust effect in deep layers in response to mispredicted compared to predictable stimuli, which cannot be ascribed to local prediction differences, as the local context is the same across conditions. This deep layer neuronal modulation to mispredicted stimuli, was not reported in a previous laminar fMRI studies contrasting expected and unexpected stimuli in the visual domain [28]. Yet, this result aligns with electrophysiological studies in animals [31,32] showing a feedforward sweep of local and global errors followed by a late decrease in the alpha/beta power band, which was indicative of a widespread global model update. In the canonical microcircuit, low frequency oscillations are associated with deep neural populations [12] and beta frequencies are thought to be negatively correlated to the BOLD response [49,50]. Thus, the increased modulation to mispredicted stimuli in deep layers, suggests that deep layers play a critical role in representing an updated internal model (see Figure 6A and 6B), which at the highest level may be stored in frontal cortical areas [51,52].

The analysis of the responses elicited by unpredictable stimuli provided further evidence to support our interpretations. We hypothesized that the presentation of unpredictable tone sequences would result in a reduced magnitude of local errors due to the rearrangement of the three contextual tones. In addition, as both the mispredicted and unpredictable sequences were presented proportionally less than the predictable sequences, we hypothesized them to both result in the updating of the internal model. In line with these hypotheses, we found no evidence for superficial and middle layer effects, yet a similar deep layer effect was present in three areas of the temporal cortex.

To study predictive processing in the temporal cortex, we used GE-BOLD, which enables efficient, sensitive imaging of the bilateral auditory cortex (Moerel et al., 2021). However, GE-BOLD is influenced by macrovascular contributions, causing spatial displacement of neuronal activity [34,54]. Other acquisition and analysis approaches have been suggested to reduce draining vein contributions in GE-BOLD [55]. For example, alternative acquisition techniques, such as spin-echo EPI [56], 3D gradient-echo and spin-echo [3D-GRASE - 57,58], and VASO [59,60], reduce macrovascular effects but suffer from limited sensitivity and coverage for auditory cortex imaging [39]. On the contrary, previous studies that did collect GE-BOLD data, used simple subtraction methods to mitigate macrovascular effects [28,61]. This assumption, however, has been shown to be inaccurate in certain applications [62]. We addressed this limitation using biophysical modeling to account for the ‘draining vein’ effect [34]. Apart from its initial validations [35,41], this model has not previously been used in neuroscientific studies. This modeling approach effectively reduced non-neuronal contributions at both middle and superficial cortical depths which are most affected by vascular draining [54], while leaving the deeper layers largely unaffected (see Figure S3 for the results obtained without the use of the model).

Interpreting our results within the canonical circuit model for PC [12] is appealing. But an alternative cortical model for PC has been recently put forward [63] and may allow a different reading of our results. In this model, and in line with previous electrophysiological research [18] prediction-errors occur in both L2/3 and L5. This suggested modified circuitry could lead to an alternative interpretation of our results, in which the activation in both the superficial and deep layers are reflective of error (local and global) computations within the cortical column and where the deep layer response we observed may not be related to global model updating. While challenging, testing these two alternative neural models (and eventually others) using non-invasive measures could be implemented within the modeling framework used here. In its current implementation, the laminar model for BOLD responses considers excitatory and inhibitory populations of neurons (recurrently connected) but separated across layers [34]. This model could be extended by considering e.g. the canonical model of PC [12] or alternative models [e.g. 63]. Model evidence, estimated within the DCM framework, would then allow adjudicating between these different neural models. Such extensions, however, may require an initial analysis of the sensitivity of the BOLD signal to the inclusion of intra-areal and intra-laminar connections.

In conclusion, the investigation of the violation of expectations across a large extent of the temporal cortex, with cortical depth dependent fMRI, revealed error-signals in superficial cortical layers, and the relevance of deep layers for broadcasting model updates. The large spatial coverage, in particular, allowed us to ground these results within the known hierarchy of temporal cortical areas - with superficial errors in belt regions feeding into middle layers in the parabelt. These findings advance our understanding of how contextual sensory information and expectations contribute to the processing of sounds and how this processing may be embedded within the human cortical (laminar) architecture.

## Methods

The data presented in this study were previously used for the evaluation of a denoising technique and published by Faes and colleagues [64]. The raw data is available on doi:10.18112/openneuro.ds004928.v1.0.0.

### 4.1 Participants

Ten participants took part in this fMRI study (5 females, mean age = 37.6 years, SD = 14.2 years) and signed informed consent before partaking in this study. None of the participants had a history of neurological or hearing disorders. Data were collected at two different locations: eight participants were scanned at the Center for Magnetic Resonance Research in Minneapolis (CMRR) and two were scanned at New York University (using identical imaging protocol with the exception of a slight difference in TR = 1650). Each local Institutional Review Board approved the experiment.

### 4.2 Experimental design and stimuli

Participants completed 6 to 8 functional runs in the scanner while they passively listened to tone sequences. These sequences were designed to manipulate predictability based on contextual information. Each sequence consisted of four sounds (see Figure 1) in which 100 ms tones were intersected with a 400 ms gap resulting in a total stimulus length of 1.6 seconds. The initial three contextual frequencies (493.9, 659.3, and 987.8 Hz) were consistently present across all conditions, though their presentation order varied depending on the specific condition (ascending, descending or scrambled). In the predictable conditions, the fourth tone adhered to the prediction generated by the three preceding sounds. On the contrary, in the ‘mispredicted’ conditions, we violated the local context by presenting a “deviant” frequency. The unpredictable conditions were designed to generate a weaker local prediction by scrambling the order of the first three frequencies (e.g. the first two tones are ascending and the last two tones were descending). With reduced predictions, we also expected the local error signal to be reduced. It’s important to note that the fourth frequency was always either 329.6 Hz or 1318.5 Hz in all conditions. The auditory stimuli were presented concomitantly with the scanner noise (i.e. no silent gap for sound presentation was used). The sequences were presented in a slow event-related design with an average inter-trial interval of 6 TRs (ranging between 5 and 7 TRs). Besides local predictability of the fourth tone, we also manipulated the global predictability by making the predictable sequences more probable (presented about 60% of the time) than the mispredicted and unpredictable sequences (each being presented about 20% of the time). Sounds were presented in the MRI scanner using MRI-compatible S14 model earbuds of the Sensimetrics Corporation (www.sens.com). During each run a fixation cross was presented in the middle of the screen.

### 4.3 MRI acquisition

At both sites, data were collected with a 7T Siemens Magnetom System with a single-channel transmit and 32-channel receive NOVA head coil (Siemens Medical Systems, Erlangen). Whole-brain anatomical T1-weighted images were collected using a Magnetization Prepared 2 Rapid Acquisition Gradient Echo (MP2RAGE) sequence [65] at a resolution of 0.75 mm isotropic (192 slices, TR = 4300 ms, TE = 2.27 ms).

The functional data were acquired with 2D gradient-echo (GE) simultaneous multi-slice (SMS)/multiband (MB) echo planar imaging (EPI) (Moeller et al., 2010; Setsompop et al., 2012) (0.8 mm isotropic, 42 slices, TR = 1600 ms, TE = 26.4 ms, MB factor 2, iPAT factor 3, 6/8 Partial Fourier, bandwidth 1190 Hz, field of view: 170 x 170 mm, matrix size: 212 x 212, phase encoding = anterior to posterior). At NYU, all acquisition parameters were the same as at CMRR with the exception of a TR of 1650 ms. In each session (except for S1), we also obtained 5 functional volumes using the opposite phase encoding (posterior to anterior).

### 4.4 Data Analysis

*Processing anatomical data.* Both the anatomical and functional data were analyzed using BrainVoyager (BV - v21.4 unless otherwise specified, Brain Innovation, Maastricht, The Netherlands). The anatomical data were upsampled to 0.4 mm isotropic, corrected for inhomogeneities, and rotated in such a way that the anterior commissure (AC) and posterior commissure (PC) were on the same plane (ACPC space). A high-resolution segmentation of the white matter (WM) and gray matter (GM) was created using a deep neural network implemented in BV (v22.0). The initial segmentation of the temporal lobes was manually corrected in ITK snap [66, v3.8]. Mid-GM surface meshes were created in BV using this corrected segmentation.

*Pre-processing functional data* consisted of slice scan time correction (sinc interpolation) and 3D motion correction (rigid body with 6 parameters) using intrasession alignment to the run closest in time to the collection of opposite phase encoding (run 1 in most participants, run 4 in two participants). Moreover, we applied temporal filtering to remove low frequencies (high-pass filtering with 7 cycles per run) and high frequencies (temporal Gaussian smoothing with a full-width half-maximum kernel of 2 data points). In two participants, we slightly diverged from the general processing pipeline (for 2 or 3 runs) because after motion correction they displayed geometric distortions (across runs). In these participants (and for these runs), we corrected for these distortions using nonlinear alignment using Advanced Normalization Tools (ANTs), prior to correcting for the distortions induced by phase encoding. In all (but one) participants, anatomical distortions introduced by phase encoding were corrected using reversed phase polarity acquisitions and Topup (FSL, v6.0.4). Finally the functional data were aligned to the upsampled anatomical data using boundary-based registration.

*Regions of interest*. The anatomical data were processed by defining five regions of interest (ROIs) on each individual’s cortical mesh surface of each hemisphere [36]. The temporal lobe was subdivided in the Heschl’s Gyrus (HG), Planum Polare (PP), Planum Temporale (PT), anterior superior temporal gyrus (aSTG) and posterior superior temporal gyrus (pSTG). These ROIs were projected back in volume space extending 3 mm inwards and outwards from the mid-GM surface. To maintain an accurate delineation of the GM ribbon, the ROIs were once more intersected with the high-resolution segmentation to identify the GM portion of each ROI. These ROIs were used as the starting point for the modeling of the hemodynamic responses (see below).

### 4.5 Modeling and statistical analysis

The CSF-gray matter and gray matter-white matter boundaries as identified in volume space by the BrainVoyager segmentation were additionally inspected with custom MATLAB (The MATHWORKS Inc., Natick, MA, USA) code and ITK-snap [66]. An equivolume approach was used to estimate the normalized voxel-wise depth estimate in the gray matter. The depth map was then downsampled to match the voxel size of the functional data (0.8 mm isotropic).

We analyzed the functional data by fitting a general linear model (GLM) in BV with the 6 conditions as predictors. For each ROI, the analysis was confined to the voxels with the strongest response to the auditory stimuli (F-statistic; p<0.05 uncorrected) at the individual subject level. The procedure we followed for modeling the responses of each ROI, layer and participant followed a methodology previously introduced [35,41]. In particular, we restricted the further modeling analysis to ROIs that contained 300 or more active voxels to ensure a sufficient number of voxels (in each depth) for model fitting. This restriction resulted in the exclusion of 4 ROIs (PP LH in one participant and aSTG LH in 3 participants). For all subjects, the event-related averages were computed per voxel and trial (of all conditions). The event-related averages were baseline corrected and comprised 7 time points. Across all subjects, ROIs and layers, outlier voxels were removed below the 0.01^st^ and above 99.99^th^ percentile (for all conditions). The event-related averages per condition were obtained by averaging over trials. Within each ROI, we identified the contribution of each voxel to several depths (7, 9, 10 and 11 bins) using normalized equally-spaced bell-shaped weighted probability maps. For every cortical depth, the hemodynamic response of two conditions was concatenated in order to use the generative model of laminar BOLD (i.e. predictable condition followed by mispredicted/unpredictable, separately for ascending and descending sequences). The responses were spaced by 20 time points randomly sampled from a white noise distribution. This was done to assure that the model fit would come back to baseline, i.e. we added an artificial ‘rest’ period to avoid overlap in modeled BOLD responses at a later stage. These added data points did not affect the modeling results (see model inversion details below). To estimate laminar neuronal activity underlying the measured BOLD responses, we used the generative model of the laminar BOLD signal [34]. This model estimates BOLD responses using a combination of venous microvasculature and ascending veins in the DCM framework. We fixed the number of neuronal depths to 3. These were then linked independently to four larger numbers of vascular depths (*K* ∈ [7, 9, 10, 11]) using neurovascular coupling. The number of vascular depths was chosen based on previous simulations [34] recommending more than six depths for the BOLD signal even if the neuronal depths are fixed to three. The model was applied per ROI and the different regions were not connected.

We generated a model space comprising eight models for each of the vascular depths. These eight models were based on the hypotheses regarding external modulatory input(s) targeting specific neuronal depths. This included the null hypothesis that responses in both conditions were the same (i.e., no influence of modulatory input was captured by the model), to modulatory input targeting only one, any two, or all three neuronal depths (superficial, middle, and deep - see Figure 3D for an illustration of the 8 models). At every layer, the model of Havlicek and Uludag [34] offers two mechanisms to modulate the response at the neuronal level, changes in the self-excitation connection (σ) or changes in the inhibitory-excitatory connection (μ). The modulation of μ has been shown to capture the changes in the post-stimulus BOLD undershoot [67]. As our experimental design did not consider long inter trial intervals hence it didn’t capture the post-stimulus undershoot. Thus, in our analyses, we limited the modulatory input to target only the self-excitation connection. The modulation of the σ parameter changes the overall amplitude and shape of the generated BOLD response by scaling excitatory-inhibitory and inhibitory-excitatory connections [see 67]. Note that this implementation of the neuronal model has no interlaminar connections and thus the modulatory inputs were modeled independently between depths.

Within this framework, the predictable condition was taken as the baseline receiving only driving input, while the second condition (mispredicted or unpredictable, modeled separately) received both driving and modulatory input. The driving input followed the structure of the input sound stimulus (4 tones, 1600 ms duration, 8 - 11.2 s inter-trial interval). The modulatory input was hypothesized to overlap (onset and duration) with driving input at the break of prediction i.e., at the fourth tone (at 1500 ms after stimulus onset) of the quartet for the mispredicted condition. For the unpredictable condition, this was modeled both at the moment at which the context was disrupted at the third tone (1000 ms after stimulus onset) and fourth tone (1500 seconds in ms stimulus onset) to capture the effect of contextual disruption on the processing of the fourth tone.

The neuronal activity at each depth was transformed into changes in cerebral blood flow (CBF), and subsequent cerebral blood volume (CBV). For each value of *K*, the simulated BOLD responses were convolved with a unique spatial blurring kernel (Gaussian point spread function, standard deviation, and scaling constant adjusted for the number of vascular depths, the curvature of the ROI, and the voxel size). We optimized neuronal and vascular parameters per vascular depth (see Table 1 for estimated parameters and prior distributions). The fixed parameters are reported in Table 2.

**Table 1.** Model parameters and priors (only the ones estimated during model inversion).

| Description |  | Prior Mean | Prior Variance |
| --- | --- | --- | --- |
| Neuronal Model |  |  |  |
| $C$ (-) | Strength of driving input to an excitatory population | [1, 1, 1] | [1, 1, 1] |
| $\sigma$ (Hz) | Excitatory self-connection | 3 | $e^{-2}$ |
| $B$ (Hz) | Modulatory effects on $\sigma$ | 0 | $e^{0.5}$ |
| Hemodynamic Model |  |  |  |
| $S_{AV}$ (-) | Slope of $CBV_0$ increase towards surface (AV) | 0 | $e^1$ |
| $\alpha_{AV}$ (-) | Exponent relating CBF to $CBV$ during steady-state (AV) | 0.35 | $e^{-5}$ |

**Table 2.** Fixed model parameters [see 35 for remaining parameters].

| Description |  | Value |
| --- | --- | --- |
| Neuronal Model |  |  |
| $\mu$ (Hz) | Inhibitory Excitatory connection | -0.8 |
| $\lambda$ (Hz) | Inhibitory gain factor | 1.8 |
| Hemodynamic Model |  |  |
| $T$ (Tesla) | Field strength | 7 |
| TR (s) | Repetition time | 1.6 |
| TE (s) | Echo time | 0.02 |
| $V_0$ (mL) | Total baseline blood volume (MV + AV) | 3 |
| $\omega_{MV}$ (-) | Fraction of total $CBV_0$ represented (MV) | 0.5 |
| $n_{MV}$ (-) | Ratio between CBF and $CMRO_2$ (MV) | 3 |

The model inversion was performed using a slightly modified version of the standard Variational Bayes approach implemented in spm12 (“spm_nlsi_GN.m”). This modification included considering only the data time points and not the white noise inserted between the data from the two conditions while performing expectation-maximization. The estimated parameters were pooled across the different values of *K* (7, 9, 10 and 11) by performing Bayesian parameter averaging (“spm_bpa.m”) for each of the eight models. After Bayesian averaging, the eight models were compared using the Free Energy metric [43]. The winning model was used to obtain the estimate of modulatory input (σ) across the three neuronal depths for each subject and ROI - representing the modulation induced by the second condition (mispredicted or unpredictable sequence) with respect to the first (predictable sequence).

To test the significance of the modulatory input across subjects and for each layer and ROI, we first computed t-tests (versus zero) and conducted non-parametric permutations for significance testing by randomly flipping (2^10^ times) the signs of the observed data and recalculating the t-statistics. Finally, we corrected for multiple tests using False Discovery Rate (FDR) across all ROIs and layers.

## Data availability

The raw data is publicly available on OPENNEURO: doi:10.18112/openneuro.ds004928.v1.0.0 [68].

## Code availability

The code used for the analysis reported in this manuscript can be found on https://github.com/lonikefaes. This will be made available upon acceptance (or earlier if requested by reviewers).

## Acknowledgements

We would like to thank Michelle Moerel for her guidance in drawing regions of interest in the auditory cortex. Moreover, we would like to thank Giancarlo Valente for his advice on the permutation tests. And finally, we would like to thank Jorie van Haren and Mahdi Enan for the useful scientific discussions. Moreover, we would like to thank the funding agencies, the National Institute of Health (grants RF1MH116978-01 and RO1DC017734-05), The European Research Council (ERC) under the European Union’s Horizon 2020 research and innovation programme (grant agreement No. 101001270), the National Science Foundation (grant 2420979) and the Biotechnology and Biological Sciences Research Council (grant BB/X013103/1, 22 UKRI-SBE).

## Author Contribution

**Lonike K Faes:** Conceptualisation, Methodology, Formal analysis, Writing - Original Draft, Writing - Review and Editing

**Isma Zulfiqar:** Conceptualisation, Methodology, Formal analysis, Writing - Original Draft, Writing - Review and Editing

**Luca Vizioli:** Investigation, Writing - Review and Editing

**Zidan Yu:** Investigation

**Yuan-Hao Wu:** Investigation

**Jiyun Shin:** Writing - Review and Editing

**Martijn A. Cloos:** Investigation, Writing - Review and Editing

**Ryszard Aukstulewicz:** Writing - Review and Editing

**Lucia Melloni:** Conceptualisation, Writing - Review and Editing, Funding Acquisition

**Kamil Uludag:** Conceptualisation, Writing - Review and Editing, Funding Acquisition

**Essa Yacoub:** Conceptualisation, Writing - Review and Editing, Funding Acquisition

**Federico De Martino:** Conceptualisation, Methodology, Supervision, Writing - Review and Editing, Funding Acquisition

## References

[1] Sohoglu E, Davis MH. Perceptual learning of degraded speech by minimizing prediction error. Proc Natl Acad Sci 2016;113. 10.1073/pnas.1523266113.

[2] Winkler I, Czigler I. Evidence from auditory and visual event-related potential (ERP) studies of deviance detection (MMN and vMMN) linking predictive coding theories and perceptual object representations. Int J Psychophysiol 2012;83:132–43. 10.1016/j.ijpsycho.2011.10.001.

[3] Winkler I, Denham SL, Nelken I. Modeling the auditory scene: predictive regularity representations and perceptual objects. Trends Cogn Sci 2009;13:532–40. 10.1016/j.tics.2009.09.003.

[4] Sedley W, Gander PE, Kumar S, Kovach CK, Oya H, Kawasaki H, et al. Neural signatures of perceptual inference. eLife 2016;5:e11476. 10.7554/eLife.11476.

[5] Haarsma J, Kok P, Browning M. The promise of layer-specific neuroimaging for testing predictive coding theories of psychosis. Schizophr Res 2022;245:68–76. 10.1016/j.schres.2020.10.009.

[6] Corlett PR, Frith CD, Fletcher PC. From drugs to deprivation: a Bayesian framework for understanding models of psychosis. Psychopharmacology (Berl) 2009;206:515–30. 10.1007/s00213-009-1561-0.

[7] Sterzer P, Adams RA, Fletcher P, Frith C, Lawrie SM, Muckli L, et al. The Predictive Coding Account of Psychosis. Biol Psychiatry 2018;84:634–43. 10.1016/j.biopsych.2018.05.015.

[8] Rao RPN, Ballard DH. Predictive coding in the visual cortex: a functional interpretation of some extra-classical receptive-field effects. Nat Neurosci 1999;2:79–87. 10.1038/4580.

[9] Friston KJ. A theory of cortical responses. Philos Trans R Soc B Biol Sci 2005;360:815–36. 10.1098/rstb.2005.1622.

[10] Felleman DJ, Van Essen DC. Distributed Hierarchical Processing in the Primate Cerebral Cortex. Cereb Cortex 1991;1:1–47. 10.1093/cercor/1.1.1.

[11] Markov NT, Ercsey-Ravasz M, Van Essen DC, Knoblauch K, Toroczkai Z, Kennedy H. Cortical High-Density Counterstream Architectures. Science 2013;342:1238406. 10.1126/science.1238406.

[12] Bastos AM, Usrey WM, Adams RA, Mangun GR, Fries P, Friston KJ. Canonical Microcircuits for Predictive Coding. Neuron 2012;76:695–711. 10.1016/j.neuron.2012.10.038.

[13] Heilbron M, Chait M. Great Expectations: Is there Evidence for Predictive Coding in Auditory Cortex? Neuroscience 2018;389:54–73. 10.1016/j.neuroscience.2017.07.061.

[14] Shipp S. Neural Elements for Predictive Coding. Front Psychol 2016;7. 10.3389/fpsyg.2016.01792.

[15] Bastos AM, Lundqvist M, Waite AS, Kopell N, Miller EK. Layer and rhythm specificity for predictive routing. Proc Natl Acad Sci 2020;117:31459–69. 10.1073/pnas.2014868117.

[16] Gallimore CG, Ricci DA, Hamm JP. Spatiotemporal dynamics across visual cortical laminae support a predictive coding framework for interpreting mismatch responses. Cereb Cortex 2023;33:9417–28. 10.1093/cercor/bhad215.

[17] Hamm JP, Shymkiv Y, Han S, Yang W, Yuste R. Cortical ensembles selective for context. Proc Natl Acad Sci 2021;118:e2026179118. 10.1073/pnas.2026179118.

[18] Jordan R, Keller GB. Opposing Influence of Top-down and Bottom-up Input on Excitatory Layer 2/3 Neurons in Mouse Primary Visual Cortex. Neuron 2020;108:1194–1206.e5. 10.1016/j.neuron.2020.09.024.

[19] Keller GB, Bonhoeffer T, Hübener M. Sensorimotor Mismatch Signals in Primary Visual Cortex of the Behaving Mouse. Neuron 2012;74:809–15. 10.1016/j.neuron.2012.03.040.

[20] Lakatos P, O’Connell MN, Barczak A, McGinnis T, Neymotin S, Schroeder CE, et al. The Thalamocortical Circuit of Auditory Mismatch Negativity. Biol Psychiatry 2020;87:770–80. 10.1016/j.biopsych.2019.10.029.

[21] Vezoli J, Magrou L, Goebel R, Wang X-J, Knoblauch K, Vinck M, et al. Cortical hierarchy, dual counterstream architecture and the importance of top-down generative networks. NeuroImage 2021;225:117479. 10.1016/j.neuroimage.2020.117479.

[22] Csercsa R, Dombovári B, Fabó D, Wittner L, Erőss L, Entz L, et al. Laminar analysis of slow wave activity in humans. Brain 2010;133:2814–29. 10.1093/brain/awq169.

[23] Halgren E, Kaestner E, Marinkovic K, Cash SS, Wang C, Schomer DL, et al. Laminar profile of spontaneous and evoked theta: Rhythmic modulation of cortical processing during word integration. Neuropsychologia 2015;76:108–24. 10.1016/j.neuropsychologia.2015.03.021.

[24] Keller CJ, Cash SS, Narayanan S, Wang C, Kuzniecky R, Carlson C, et al. Intracranial microprobe for evaluating neuro-hemodynamic coupling in unanesthetized human neocortex. J Neurosci Methods 2009;179:208–18. 10.1016/j.jneumeth.2009.01.036.

[25] Kok P, Bains LJ, van Mourik T, Norris DG, de Lange FP. Selective Activation of the Deep Layers of the Human Primary Visual Cortex by Top-Down Feedback. Curr Biol 2016;26:371–6. 10.1016/j.cub.2015.12.038.

[26] Muckli L, De Martino F, Vizioli L, Petro LS, Smith FW, Ugurbil K, et al. Contextual Feedback to Superficial Layers of V1. Curr Biol 2015;25:2690–5. 10.1016/j.cub.2015.08.057.

[27] Aitken F, Menelaou G, Warrington O, Koolschijn RS, Corbin N, Callaghan MF, et al. Prior expectations evoke stimulus-specific activity in the deep layers of the primary visual cortex. PLOS Biol 2020;18:e3001023. 10.1371/journal.pbio.3001023.

[28] Thomas ER, Haarsma J, Nicholson J, Yon D, Kok P, Press C. Predictions and errors are distinctly represented across V1 layers. Curr Biol 2024;34:2265-2271.e4. 10.1016/j.cub.2024.04.036.

[29] Yu Y, Huber L, Yang J, Jangraw DC, Handwerker DA, Molfese PJ, et al. Layer-specific activation of sensory input and predictive feedback in the human primary somatosensory cortex. Sci Adv 2019;5:eaav9053. 10.1126/sciadv.aav9053.

[30] Parras GG, Nieto-Diego J, Carbajal GV, Valdés-Baizabal C, Escera C, Malmierca MS. Neurons along the auditory pathway exhibit a hierarchical organization of prediction error. Nat Commun 2017;8:2148. 10.1038/s41467-017-02038-6.

[31] Chao ZC, Takaura K, Wang L, Fujii N, Dehaene S. Large-Scale Cortical Networks for Hierarchical Prediction and Prediction Error in the Primate Brain. Neuron 2018;100:1252–1266.e3. 10.1016/j.neuron.2018.10.004.

[32] Jiang Y, Komatsu M, Chen Y, Xie R, Zhang K, Xia Y, et al. Constructing the hierarchy of predictive auditory sequences in the marmoset brain. eLife 2022;11:e74653. 10.7554/eLife.74653.

[33] Dürschmid S, Edwards E, Reichert C, Dewar C, Hinrichs H, Heinze H-J, et al. Hierarchy of prediction errors for auditory events in human temporal and frontal cortex. Proc Natl Acad Sci 2016;113:6755–60. 10.1073/pnas.1525030113.

[34] Havlicek M, Uludağ K. A dynamical model of the laminar BOLD response. NeuroImage 2020;204:116209. 10.1016/j.neuroimage.2019.116209.

[35] Uludag K, Havlicek M. Determining laminar neuronal activity from BOLD fMRI using a generative model. Prog Neurobiol 2021;207:102055. 10.1016/j.pneurobio.2021.102055.

[36] Kim J, Crespo-Facorro B, Andreasen NC, O’Leary DS, Zhang B, Harris G, et al. An MRI-Based Parcellation Method for the Temporal Lobe 2000.

[37] De Martino F, Esposito F, van de Moortele P-F, Harel N, Formisano E, Goebel R, et al. Whole brain high-resolution functional imaging at ultra high magnetic fields: an application to the analysis of resting state networks. NeuroImage 2011;57:1031–44. 10.1016/j.neuroimage.2011.05.008.

[38] Moeller S, Yacoub E, Olman CA, Auerbach E, Strupp J, Harel N, et al. Multiband multislice GE-EPI at 7 tesla, with 16-fold acceleration using partial parallel imaging with application to high spatial and temporal whole-brain fMRI. Magn Reson Med 2010;63:1144–53. 10.1002/mrm.22361.

[39] Moerel M, Yacoub E, Gulban OF, Lage-Castellanos A, De Martino F. Using high spatial resolution fMRI to understand representation in the auditory network. Prog Neurobiol 2021;207:101887. 10.1016/j.pneurobio.2020.101887.

[40] Turner R. How Much Cortex Can a Vein Drain? Downstream Dilution of Activation-Related Cerebral Blood Oxygenation Changes. NeuroImage 2002;16:1062–7. 10.1006/nimg.2002.1082.

[41] Kashyap S, Ivanov D, Havlicek M, Huber L, Poser BA, Uludağ K. Sub-millimetre resolution laminar fMRI using Arterial Spin Labelling in humans at 7 T. PLOS ONE 2021;16:e0250504. 10.1371/journal.pone.0250504.

[42] Auksztulewicz R, Schwiedrzik CM, Thesen T, Doyle W, Devinsky O, Nobre AC, et al. Not All Predictions Are Equal: “What” and “When” Predictions Modulate Activity in Auditory Cortex through Different Mechanisms. J Neurosci 2018;38:8680–93. 10.1523/JNEUROSCI.0369-18.2018.

[43] Friston KJ, Stephan KE. Free-energy and the brain. Synthese 2007;159:417–58. 10.1007/s11229-007-9237-y.

[44] Moerel M, De Martino F, Formisano E. An anatomical and functional topography of human auditory cortical areas. Front Neurosci 2014;8. 10.3389/fnins.2014.00225.

[45] Hackett TA, Stepniewska I, Kaas JH. Subdivisions of auditory cortex and ipsilateral cortical connections of the parabelt auditory cortex in macaque monkeys. J Comp Neurol 1998;394:475–95. 10.1002/(SICI)1096-9861(19980518)394:4<475::AID-CNE6>3.0.CO;2-Z.

[46] Hackett TA, Preuss TM, Kaas JH. Architectonic identification of the core region in auditory cortex of macaques, chimpanzees, and humans. J Comp Neurol 2001;441:197–222. 10.1002/cne.1407.

[47] Rauschecker JP, Tian B. Processing of Band-Passed Noise in the Lateral Auditory Belt Cortex of the Rhesus Monkey. J Neurophysiol 2004;91:2578–89. 10.1152/jn.00834.2003.

[48] Romanski LM, Averbeck BB. The Primate Cortical Auditory System and Neural Representation of Conspecific Vocalizations. Annu Rev Neurosci 2009;32:315–46. 10.1146/annurev.neuro.051508.135431.

[49] Conner CR, Ellmore TM, Pieters TA, DiSano MA, Tandon N. Variability of the Relationship between Electrophysiology and BOLD-fMRI across Cortical Regions in Humans. J Neurosci 2011;31:12855–65. 10.1523/JNEUROSCI.1457-11.2011.

[50] Scheeringa R, Fries P, Petersson K-M, Oostenveld R, Grothe I, Norris DG, et al. Neuronal Dynamics Underlying High- and Low-Frequency EEG Oscillations Contribute Independently to the Human BOLD Signal. Neuron 2011;69:572–83. 10.1016/j.neuron.2010.11.044.

[51] Alexander WH, Brown JW. Frontal cortex function as derived from hierarchical predictive coding. Sci Rep 2018;8:3843. 10.1038/s41598-018-21407-9.

[52] Bellet ME, Gay M, Bellet J, Jarraya B, Dehaene S, van Kerkoerle T, et al. Spontaneously emerging internal models of visual sequences combine abstract and event-specific information in the prefrontal cortex. Cell Rep 2024;43:113952. 10.1016/j.celrep.2024.113952.

[53] Hackett TA. Organization and Correspondence of the Auditory Cortex of Humans and Nonhuman Primates. Evol. Nerv. Syst., Elsevier; 2007, p. 109–19. 10.1016/B0-12-370878-8/00012-4.

[54] Markuerkiaga I, Barth M, Norris DG. A cortical vascular model for examining the specificity of the laminar BOLD signal. NeuroImage 2016;132:491–8. 10.1016/j.neuroimage.2016.02.073.

[55] Yacoub E, Shmuel A, Logothetis N, Uğurbil K. Robust detection of ocular dominance columns in humans using Hahn Spin Echo BOLD functional MRI at 7 Tesla. NeuroImage 2007;37:1161–77. 10.1016/j.neuroimage.2007.05.020.

[56] Yacoub E, Duong TQ, Van De Moortele P-F, Lindquist M, Adriany G, Kim S-G, et al. Spin-echo fMRI in humans using high spatial resolutions and high magnetic fields. Magn Reson Med 2003;49:655–64. 10.1002/mrm.10433.

[57] Feinberg DA, Harel N, Ramanna S, Ugurbil K, Yacoub E. Submillimeter single-shot 3D GRASE with inner volume selection for T2-weighted fMRI applications at 7T. Proc Int Soc Mag Reson Med 2008;16:2373.

[58] Oshio K, Feinberg DA. GRASE (Gradient-and Spin-Echo) imaging: A novel fast MRI technique. Magn Reson Med 1991;20:344–9. 10.1002/mrm.1910200219.

[59] Huber L, Ivanov D, Krieger SN, Streicher MN, Mildner T, Poser BA, et al. Slab-selective, BOLD-corrected VASO at 7 Tesla provides measures of cerebral blood volume reactivity with high signal-to-noise ratio: SS-SI-VASO Measures Changes of CBV in Brain. Magn Reson Med 2014;72:137–48. 10.1002/mrm.24916.

[60] Lu H, Golay X, Pekar JJ, van Zijl PCM. Functional magnetic resonance imaging based on changes in vascular space occupancy. Magn Reson Med 2003;50:263–74. 10.1002/mrm.10519.

[61] Heynckes M, Lage-Castellanos A, De Weerd P, Formisano E, De Martino F. Layer-specific correlates of detected and undetected auditory targets during attention. Curr Res Neurobiol 2023;4:100075. 10.1016/j.crneur.2023.100075.

[62] Shmuel A, Chaimow D, Raddatz G, Ugurbil K, Yacoub E. Mechanisms underlying decoding at 7 T: ocular dominance columns, broad structures, and macroscopic blood vessels in V1 convey information on the stimulated eye. NeuroImage 2010;49:1957–64. 10.1016/j.neuroimage.2009.08.040.

[63] Nejad KK, Anastasiades P, Hertäg L, Costa RP. Self-supervised predictive learning accounts for layer-specific cortical observations. BioRxiv 2024.

[64] Faes LK, Lage-Castellanos A, Valente G, Yu Z, Cloos MA, Vizioli L, et al. Evaluating the effect of denoising submillimeter auditory fMRI data with NORDIC. Imaging Neurosci 2024;2:1–18. 10.1162/imag_a_00270.

[65] Marques JP, Kober T, Krueger G, van der Zwaag W, Van de Moortele P-F, Gruetter R. MP2RAGE, a self bias-field corrected sequence for improved segmentation and T1-mapping at high field. NeuroImage 2010;49:1271–81. 10.1016/j.neuroimage.2009.10.002.

[66] Yushkevich PA, Piven J, Hazlett HC, Smith RG, Ho S, Gee JC, et al. User-guided 3D active contour segmentation of anatomical structures: Significantly improved efficiency and reliability. NeuroImage 2006;31:1116–28. 10.1016/j.neuroimage.2006.01.015.

[67] Havlicek M, Roebroeck A, Friston K, Gardumi A, Ivanov D, Uludag K. Physiologically informed dynamic causal modeling of fMRI data. NeuroImage 2015;122:355–72. 10.1016/j.neuroimage.2015.07.078.

[68] Faes LK, Lage-Castellanos A, Valente G, Yu Z, Cloos MA, Vizioli L, et al. Predictive Processing in the Auditory Cortex measured at 7T 2024. doi:10.18112/openneuro.ds004928.v1.0.0.

