## Supplementary Material for "Predictive acoustical processing in human cortical layers"

### Supplementary Materials

#### Data preprocessing

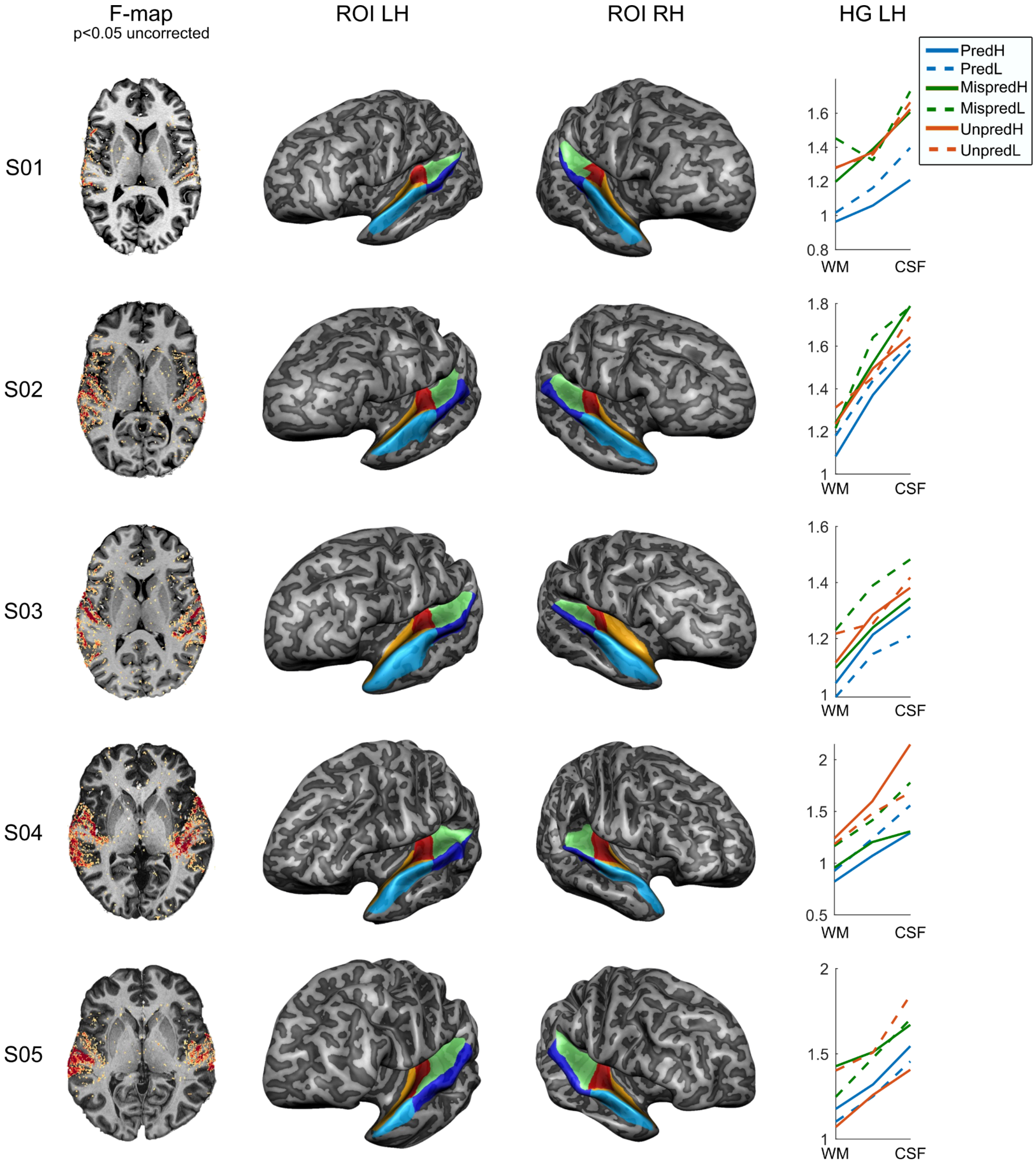


Figure continues on the next page

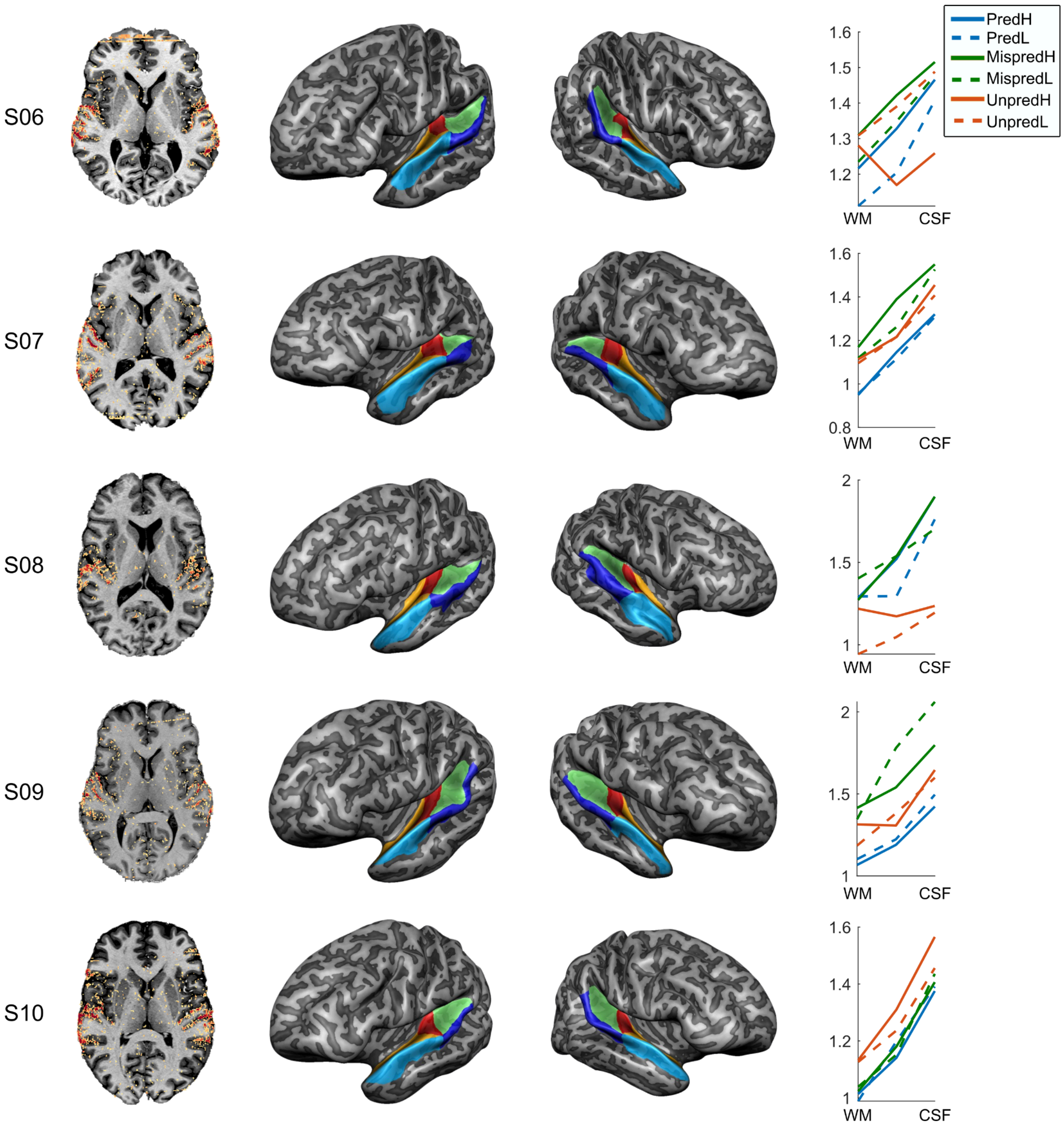


**Figure S1. Data before modeling.** The left column illustrates the active voxels responding to sounds at qFDR<0.05 uncorrected. These voxels (confined to the ROIs) were selected to be included in the analysis. The middle two columns show the parcellation of ROIs on a mid-GM surface (HG = red, PP = orange, PT = green, aSTG = light blue, pSTG = dark blue). The rightmost column illustrates the draining effect for all six conditions (blue = predictable, orange = mispredicted, yellow = unpredictable).

#### Best Model

Figure S2A shows once more the eight models and their respective target of the modulatory input (same as in Figure 3D). In each hemisphere, ROI and participant, we determined the best model. On average, the winning model (best explaining the observed fMRI data) was the model that had modulatory input in all layers (model 4), illustrated for both hemispheres in Figure S2B. We then used the modulation values for the best model (in that specific hemisphere, ROI and subject) for further analysis.

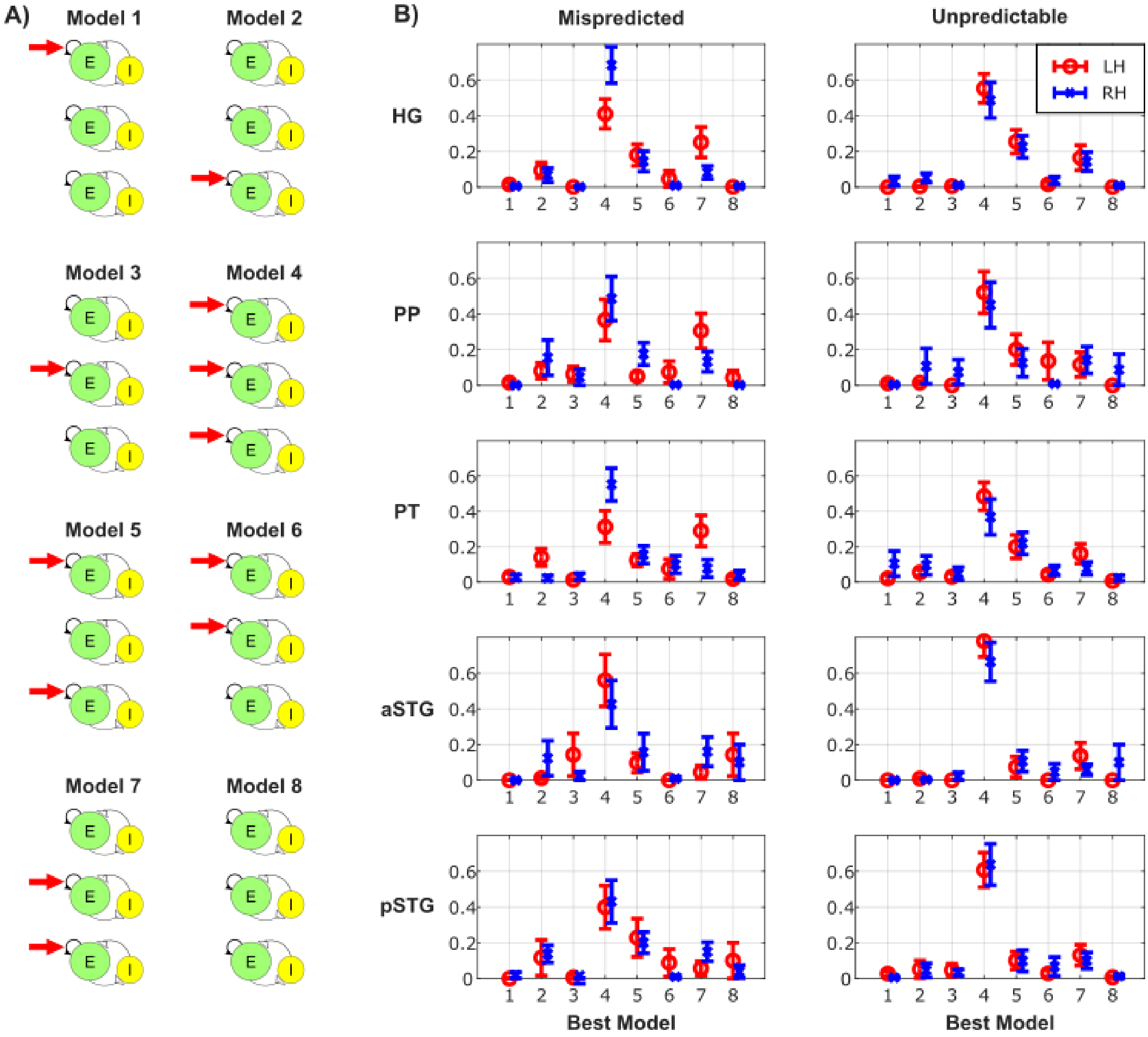


**Figure S2. Best model.** A) Illustration of the model space including a specification of which number corresponds to which model (same as Figure 3D). B) Histograms illustrating the winning model for both hemispheres and all ROIs indicating the spread across subjects. The error bars illustrate the standard error. On average model 4 was the best model, indicating modulation in all three layers.

**Data without modeling**

Prior to modeling we examined the nature of the estimated BOLD responses. To do so, we considered the maximum response (one TR) in the event-related average of the individual conditions and averaged across trails within each layer (3 depths - to compare to the modeling results) and ROI. As illustrated in Figure S3A, the anticipated increase in GE-BOLD towards the surface is evident for each ROI and each condition. Note that this figure illustrates the average across participants (in Figures S1 right column, we plot the results for single individuals).

Figure S3B depicts both the modeling and original data results for the difference between the mispredicted and predictable stimuli (top), and the unpredictable and predictable stimuli (bottom). To compare the two respective conditions in the data without modeling, we have taken the maximum response of the event-related averages to remain close to the information that the model receives as input. However, it is important to note that the model takes the full event-related average into account instead of only the maximum value. As expected, the general tendency for the model is to reduce the effects in superficial and middle layers as a part of this effect is explained by vascular draining instead of responses at the neural level [54].

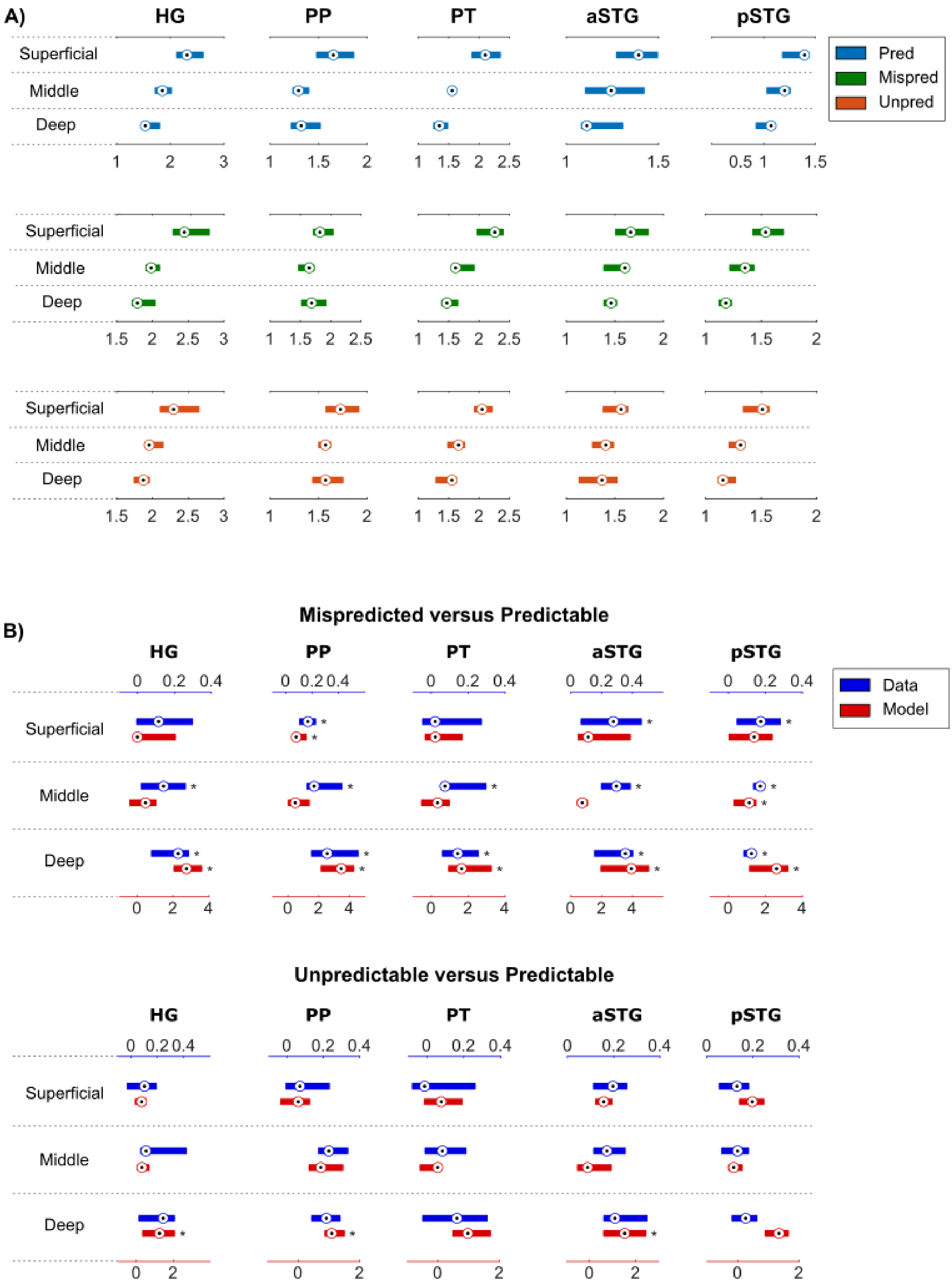


**Figure S3. Data without modeling.** A) An increase towards the surface is visible in the data without modeling for all three conditions. B) Data with and without modeling plotted in the same figure for the mispredicted condition (top) and unpredictable condition (bottom). Data without modeling represents the difference in maximum percent signal change between the mispredicted/unpredictable condition and predictable conditions. The modeling data is the same as in Figures 4 and 5 of the main manuscript. The x-axis at the top refers to the difference in maximum response in the data without modeling and the x-axis at the bottom refers to the modulation. * indicates qFDR <0.05.

#### Statistical results

#### Table S1. All statistical values of mispredicted versus predictable

| | HG | | PP | | PT | | aSTG | | pSTG | |
|---|---|---|---|---|---|---|---|---|---|---|
| | t(9) | $p_{FDR-corr}$ | t(9) | $p_{FDR-corr}$ | t(9) | $p_{FDR-corr}$ | t(9) | $p_{FDR-corr}$ | t(9) | $p_{FDR-corr}$ |
| Superficial | 1.75 | 0.166 | 3.75 | 0.005 | 1.31 | 0.226 | 2.24 | 0.100 | 2.94 | 0.051 |
| Middle | 1.70 | 0.166 | 1.49 | 0.212 | 0.86 | 0.422 | 2.47 | 0.055 | 4.69 | 0.005 |
| Deep | 4.88 | 0.008 | 6.53 | 0.005 | 4.37 | 0.005 | 6.90 | 0.005 | 5.88 | 0.005 |

#### Table S2. All statistical values of unpredictable versus predictable

| | HG | | PP | | PT | | aSTG | | pSTG | |
|---|---|---|---|---|---|---|---|---|---|---|
| | t(9) | $p_{FDR-corr}$ | t(9) | $p_{FDR-corr}$ | t(9) | $p_{FDR-corr}$ | t(9) | $p_{FDR-corr}$ | t(9) | $p_{FDR-corr}$ |
| Superficial | 0.00 | 1.000 | -0.63 | 0.938 | 0.53 | 0.938 | 0.85 | 0.896 | 0.06 | 1.000 |
| Middle | 0.21 | 1.000 | 1.54 | 0.391 | -0.57 | 0.936 | -0.02 | 1.000 | 0.22 | 1.000 |
| Deep | 3.41 | 0.049 | 3.57 | 0.049 | 3.20 | 0.059 | 4.70 | 0.049 | 1.61 | 0.393 |
